# Systemic inflammation modulates the levels of EBI2 and 7α,25-OHC synthesizing (CH25H, CYP7B1) and degrading (HSD3B7) enzymes in brain microvascular cells

**DOI:** 10.1101/2023.04.16.537063

**Authors:** Fionä Caratis, Bartosz Karaszewski, Ilona Klejbor, Tomomi Furihata, Aleksandra Rutkowska

**Affiliations:** Brain Diseases Centre, Medical University of Gdansk, Gdansk, Poland; Department of Biochemistry, Medical University of Gdansk, Gdansk, Poland; Department of Adult Neurology, Medical University of Gdansk & University Clinical Center, Gdansk, Poland; Department of Anatomy, Collegium Medicum, Jan Kochanowski University in Kielce, Kielce, Poland; Laboratory of Clinical Pharmacy and Experimental Therapeutics, School of Pharmacy, Tokyo University of Pharmacy and Life Sciences, Hachioji, Tokyo, Japan; Department of Anatomy and Neurobiology, Medical University of Gdansk, Gdansk, Poland

**Keywords:** neuroinflammation, oxysterols, EBI2, blood-brain barrier, microvessels

## Abstract

The endogenous ligand for the EBI2 receptor, oxysterol 7α,25OHC, crucial for immune responses, is finely regulated by CH25H, CYP7B1 and HSD3B7 enzymes. Lymphoid stromal cells and follicular dendritic cells within T cell follicles maintain a gradient of 7α,25OHC, with stromal cells increasing and dendritic cells decreasing its concentration. This gradient is pivotal for proper B cell positioning in lymphoid tissue. In the animal model of multiple sclerosis, the experimental autoimmune encephalomyelitis, the levels of 7α,25OHC rapidly increase in the central nervous system driving the migration of EBI2 expressing immune cells through the blood-brain barrier (BBB). To explore if blood vessel cells in the brain express these enzymes, we examined normal mouse brain microvessels and studied their expression changes during inflammation. EBI2 was abundantly expressed in endothelial cells, pericytes/smooth muscle cells, and astrocytic endfeet. CH25H, CYP7B1, and HSD3B7 were variably detected in each cell type, suggesting their active involvement in oxysterol 7α,25OHC synthesis and gradient maintenance under normal conditions. Under acute inflammatory conditions, EBI2 and synthesizing enzyme modulation occurred in brain vasculature, with variations based on the enzyme and cell type. Significant species-specific differences emerged in EBI2 and enzyme levels between mouse and human BBB-forming cells. Overall, our investigation suggest a direct role of the brain microvascular cells in regulating 7α,25OHC levels, shedding light on potential therapeutic targets for neuroinflammatory disorders.

## INTRODUCTION

The GPR183 (also known as Epstein-Barr virus-induced gene 2 (EBI2)) is a G protein-coupled receptor discovered in 1993 in EBV-infected B lymphocytes (Birkenbach et al. 1993). Since then, EBI2 was shown to play important roles in the regulation of the innate and adaptive immune system, cellular migration (Pereira et al. 2009; Gatto et al. 2009; Hannedouche et al. 2011; Preuss et al. 2014; Rutkowska et al. 2015, 2016), inflammatory signalling (Clottu et al. 2017; Wanke et al. 2017; Rutkowska et al. 2018) and myelination in the central nervous system (CNS) (Nelissen et al. 2012; Rutkowska et al. 2017; Velasco-Estevez et al. 2021). Its most potent endogenous agonist is oxysterol 7α,25-dihydroxycholesterol (7α,25OHC) (Hannedouche et al. 2011; Liu et al. 2011). 7α,25OHC is synthesized from cholesterol by the sequential enzymatic activity of cholesterol 25-hydroxylase (CH25H) and 25-hydroxycholesterol 7-alpha-hydroxylase (CYP7B1) and metabolized by 3beta-hydroxy-Delta(5)-C27-steroid oxidoreductase (HSD3B7) (Russell 2003) **(Figure 1A)**. 7α,25OHC was repeatedly shown to direct immune cell migration via an oxysterol gradient, both *in vivo* and *in vitro*, in EBI2-expressing cells demonstrating its key regulatory role in immune cell function and immunity (Hannedouche et al. 2011; Elain et al. 2014; Gessier et al. 2014; Rutkowska et al. 2016; Clottu et al. 2017).

**Fig. 1.**
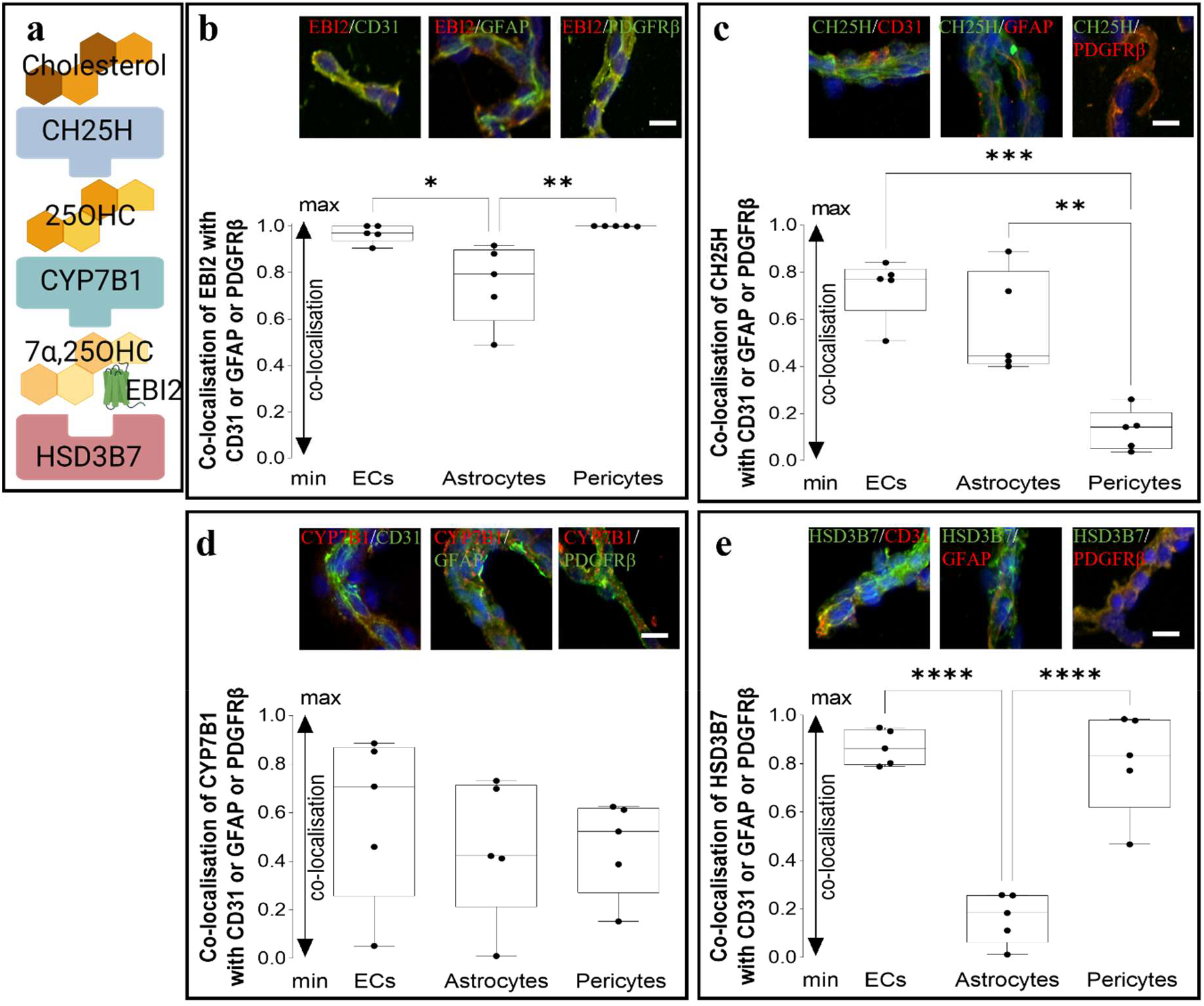
The oxysterol 7α,25OHC synthesising (CH25H, CYP7B1) and degrading (HSD3B7) enzymes are present in isolated mouse brain microvessels. a. Schematic representation of the synthetic and degrading pathway of 7α,25OHC, the endogenous ligand for EBI2. b. EBI2 highly co-colocalised with pericytes (PDGFRβ+ 0.99 +/- 0.003) and ECs (CD31+ 0.97 +/-0.09). c. CH25H was detected in ECs (CD31+), the astrocytic endfeet (GFAP+) and pericytes (PDGFRβ+) with highest co-localisation with the EC marker CD31 (0.73 +/- 0.33) and lowest with the pericyte marker PDGFRβ (0.13 +/- 0.22). d. CYP7B1 co-localised with CD31 (0.59 +/-0.83), GFAP (0.45 +/- 0.72) and PDGFRβ (0.46 +/-0.47). e. HSD3B7 mainly co-localised with ECs (0.87 +/-0.16) and pericytes (0.81 +/-0.52) with low co-localisation with astrocytes (0.16 +/-0.52). Data shown as box and whiskers with median (solid line); 1=complete co-localisation, 0=no co-localisation, one-way ANOVA and Tukey’s multiple comparisons test, *p<0.05, **p<0.01, ***p<0.001, ****p<0.0001 vs. corresponding vehicle, N=microvessels isolated from 8 mouse brains (combined). Scale 10 μm.

In normal CNS, the entry of immune cells is limited by the blood-brain barrier (BBB), a physical barrier established at the level of the brain microvascular endothelial cells (ECs) by tight junction proteins and further supported and regulated by pericytes and astrocytic endfeet, among other cells (Armulik et al. 2010). The function of the barrier is compromised by a range of factors including inflammation, neurodegeneration, stroke and brain trauma, to name just a few. In multiple sclerosis (MS), a chronic inflammatory disease of the CNS, disruption of the BBB occurs focally and at very early stages facilitating the entry of peripherally activated lymphocytes into the brain parenchyma. In the animal model of MS, the experimental autoimmune encephalomyelitis (EAE), increased concentration of 7α,25OHC in mouse CNS was independently reported by two groups (Wanke et al. 2017; Mutemberezi et al. 2018). Wanke and colleagues (Wanke et al. 2017) established that the increase in 7α,25OHC levels resulted from the upregulation of CH25H by microglia and CYP7B1 by infiltrating lymphocytes during simultaneous downregulation of HSD3B7. Importantly, the increased concentration of the EBI2 ligand enhanced the migration of autoreactive T cells into the CNS during the early phases of transfer EAE, exacerbating the disease course. Similar conclusions were drawn in a study using the EAE model and CH25H knock-out mice (Chalmin et al. 2015). The EAE severity was significantly attenuated in the CH25H deficient mice in part by reduced trafficking of encephalitogenic CD4+ T cells into the CNS. In a more recent study, the same group demonstrated that knock-out of CH25H specifically in ECs attenuates the course of EAE (Ruiz et al. 2023).

The levels of oxysterols in the plasma and the cerebrospinal fluid (CSF) also change dynamically in humans in the course of inflammatory diseases. Crick and colleagues (2017) measured a wide range of oxysterols in the CSF and plasma of patients diagnosed with relapsing-remitting MS, clinically isolated syndrome (CIS), neurodegenerative diseases including Parkinson’s and Alzheimer’s disease, amyotrophic lateral sclerosis and other inflammatory CNS diseases. Specifically in MS patients, the levels of 25OHC, the precursor to 7α,25OHC, were reduced in the plasma and increased only in the CSF in relapsing-remitting MS indicating disease-specific modulation of oxysterol levels in the CNS.

Apart from the recent identification of CH25H as a synthesizing enzyme by CNS ECs (Chalmin et al. 2015), the expression of EBI2 and the specific enzymes involved in the synthesis (CH25H, CYP7B1) and degradation (HSD3B7) of 7α,25OHC in the brain microvascular cells have not been investigated (Ruiz et al. 2023). Here, we examine whether EBI2 and the enzymes in the 7α,25OHC synthetic pathway (CH25H, CYP7B1, HSD3B7) are differentially expressed in the brain vascular cells and if systemic inflammation modulates their levels specifically in these cells. Regulation of 7α,25OHC levels by the brain microvascular cells may help in the identification of novel drug targets for the modulation of neuroinflammatory signalling and immune cell trafficking into the CNS.

## MATERIALS AND METHODS

### Animals

All animal experiments were approved by the Local Ethical Committee for Animal Experiments in Bydgoszcz, Poland under licence numbers 27/2019 and 38/2021. The C57BL/6 male mice were housed in standard cages with an enriched environment under 12-hour day and night cycles. The air in the room was exchanged 15 times per hour, the temperature was kept at 20-23°C and the humidity was between 50-60%. The animals had unrestricted access to food and water.

### LPS *in vivo* model

Forty-eight three-month-old male C57BL/6 mice were used. The mice received intraperitoneal (i.p.) injections of 0,9% NaCl (vehicle) or 2 mg/kg LPS for 12 or 24 hours (h). The mice were anaesthetised with isoflurane and perfused with NaCl for subsequent biochemical analysis or 2% paraformaldehyde (PFA) for immunohistochemistry. Whole brains were removed, snap-frozen and stored at -80°C for subsequent biochemical analysis or put into the O.C.T. compound and cut into 40 μm sections with a cryostat. The cut slices were submerged in anti-freeze and stored at -20°C until used for immunohistochemistry. The snap-frozen brains (right hemispheres) were homogenised with a pestle and a mortar in liquid nitrogen. The resulting powder was then suspended in 400 μL of fenozol/50 mg of tissue and frozen at -80°C until needed. The left hemispheres were homogenised with a Dounce homogeniser in RIPA buffer supplemented with a protein inhibitor cocktail (700 μl/100 mg of tissue) and centrifuged at 13.4k rpm for 20 minutes at 4°C. The supernatant was then aliquoted for ELISAs and stored at -80°C until needed. The pellet was resuspended in 2% SDS and sonicated with a bandelin sonicator (1.5 M) four times for 10 seconds at an amplitude of 75% and one final time for 10 seconds at 85% amplitude. Samples were then centrifuged at 13.4 k rpm for 20 minutes at 4°C and the supernatant was aliquoted for western blot (WB) and stored at -80°C until needed.

### Mouse microvessel isolation

Postnatal day 10 (P10) C57BL/6 mice were used for brain microvessel isolation according to the protocols by Paraiso et al. and Hartz et al. (Hartz et al. 2018; Paraiso et al. 2020). Briefly, the brain was extracted from the skull, the cerebral cortex was separated from the rest of the brain and the meninges were removed. Cortices were placed in the isolation buffer: 5 mM glutaMAX and 1 mM sodium pyruvate in DPBS with calcium and magnesium, pH 7.4 at 4°C. Cortices were then homogenised in the isolation buffer with a Dounce homogeniser for 20 strokes. The homogenate was moved to 2 ml tubes and then 40% polysucrose 400 (P7798-100 g, Merck) in DPBS with calcium and magnesium was added in equal volumes for a final concentration of 20%. After vigorous shaking, the homogenates were centrifuged at 7.7k rpm for 15-20 minutes at 4°C (fixed angle rotor). The pellet was resuspended in 1% bovine serum albumin (BSA) in the isolation buffer and was then filtered through a 300 μm mesh. The mesh was washed with 4 ml of the BSA-containing isolation buffer. The flow-through was filtered through a 30 μm cell strainer twice and the mesh was washed twice with BSA-containing isolation buffer before discarding the filtrate. The filter was turned upside-down and washed twice with BSA-isolation buffer to wash off the capillaries. After collection, the capillaries were centrifuged at 1500 rpm for 3 minutes at 4°C and resuspended in the isolation buffer. The capillaries were centrifuged and resuspended in the isolation buffer for subsequent analysis. The capillaries were left to adhere to the bottom of the plate in the isolation buffer in the incubator at 37°C and 5% CO_2_.

### Real-time quantitative polymerase chain reaction (RT-qPCR)

RNA was isolated using the total RNA mini plus kit (036-100, AA Biotech, Poland) according to the manufacturer’s protocol. The cDNA synthesis was performed with a transcriba kit (4000, AA Biotech, Poland) using the following programme: 60 minutes at 42°C followed by 5 minutes at 70°C or using the high-capacity cDNA reverse transcription kit (4368814, ThermoFisher) with the following programme: 10 minutes at 25°C followed by 120 minutes at 37°C and 5 seconds at 85°C. The RT-qPCR was then performed with the TaqMan fast advanced master mix (4444557, ThermoFisher) or the Sensitive RT HS-PCR Mix (2017-2000, AA Biotech, Gdansk, Poland) on the LightCycler480 (Roche) according to the manufacturer’s protocol. All samples were run in duplicate. The following FAM dye-labelled Taqman (Applied Biosystems) primers were used: β-actin (Mm02619580_g1), GAPDH (Mm99999915_g1), EBI2 (Mm02620906_s1), CH25H (Mm00515486_s1), CYP7B1 Mm00484157_m1), HSD3B7 (Mm01159156_g1), Occludin (Mm00500912_m1), N-cadherin (Mm01162490_m1). The relative gene expression was determined after normalisation to the housekeeping gene using the ΔCt (baseline gene expression in normal tissue) or ΔΔCt for comparisons of gene expression between treated and untreated samples.

### Immunohistochemistry of mouse brain sections

The mouse sections in anti-freeze were washed in PBS for 3 x 5 minutes and then incubated in cold 4% PFA for 5 minutes at room temperature (RT) followed by incubation in 20% ice-cold methanol for 1 minute. The sections were then washed for 3 x 5 minutes with PBS and blocked for 6 hours at RT with PBS supplemented with 10% BSA, 0.5% Triton-X and 1% normal goat serum (NGS). Blocked sections were incubated overnight at RT with primary antibodies diluted in the following solution: PBS supplemented with 2% BSA and 0.1% Triton-X. Only validated and previously published primary antibodies from established suppliers were used in the study. Partial validation was additionally performed for this study by performing immunohistochemistry with isotype control antibodies and using EBI2 knock-out (KO) brain sections **(Supplemental Figure 1)**. Primary antibodies (1:100 unless stated otherwise) used were: goat anti-EBI2 (RRID: AB_1090369, ab121001, Abcam,), rabbit polyclonal CH25H (600-401-MM8, ThermoFisher), mouse monoclonal CYP7B1 (OTI1G7) (TA807549, ThermoFisher), rabbit polyclonal HSD3B7 (RRID: AB_10856786, BS-2366R, ThermoFisher), mouse monoclonal Iba1 (GT10312) (1:200) (RRID: AB_2735228, MA5-27726, ThermoFisher), rabbit Iba1 (RRID:AB_839504, 019-19741, Wako), mouse monoclonal GFAP antibody (1:200) (RRID: AB_2827276, SAB5201104, Sigma-Aldrich), rabbit monoclonal GFAP (1:200) (RRID: AB_2631098, 12389, CellSignalling), mouse monoclonal CD31 (PECAM-1) (RRID: AB_10596359, BMS137, eBioscience), rabbit polyclonal CD31 (PECAM-1) (RRID: AB_10981955, PA5-16301, ThermoFisher), rabbit monoclonal PDGFRβ (RRID: AB_10985851, MA5-15143, ThermoFisher), goat polyclonal PDGFRb (RRID: AB_2162633, AF1042, R&D Systems), rabbit polyclonal occludin (RRID: AB_2533468, 40-4700, ThermoFisher), mouse monoclonal N-cadherin (3B9) (RRID: AB_2313779, 333900, ThermoFisher). The following day, sections were washed 3 x 10 minutes in PBS and incubated for 2 hours at RT with the following secondary antibodies (1:500): goat anti-mouse IgG (H+L) cross-adsorbed Alexa Fluor 546 (RRID: AB_2534071, A-11003, ThermoFisher), goat anti-mouse IgG (H+L) cross-adsorbed Alexa Fluor 488 (RRID: AB_2338840, 115-545-003-20, JacksonImmuno), goat anti-rabbit IgG (H+L) cross-adsorbed Alexa Fluor 488 (RRID: AB_2630356, ab150077, Abcam), donkey anti-goat IgG (H+L) cross-adsorbed Alexa Fluor 546 (RRID: AB_2534103, A11056, ThermoFisher)and Hoechst33342 (H1399, ThermoFisher). The sections were then washed 3 x 10 minutes in PBS, air-dried, mounted with Keiser’s gelatine and imaged using a confocal Zeiss LSM880 microscope.

### Immunohistochemistry of isolated brain microvessels

After isolation, mouse capillaries were plated in PBS on poly-D-lysine coated 8-well μ-slide plates (80826, Ibidi) and left to adhere for 2 hours in the incubator at 37°C and 5% CO_2_. The media was removed and the capillaries were left to air dry for 30 minutes. The capillaries were washed once in PBS and then fixed in 4% PFA for 10 minutes. The PFA was removed and 50 μl of ice-cold methanol was added for 1 minute. The capillaries were then washed twice with PBS at RT and blocked for 1 hour in 0.5% NGS, 1% BSA and 0.1% Tween20 in PBS. Subsequently, the capillaries were incubated overnight at 4°C in 0.5% BSA and 0.05% Tween20 in PBS with the primary antibodies. Primary antibodies used for mouse microvessels (1:100) were: goat polyclonal EBI2 (RRID: AB_10903697, ab121001, Abcam), rabbit polyclonal CH25H (600-401-MM8, ThermoFisher), mouse monoclonal CYP7B1 (OTI1G7) (TA807549, ThermoFisher), rabbit polyclonal HSD3B7 (RRID: AB_10856786, BS-2366R, ThermoFisher), rabbit monoclonal GFAP (RRID: AB_2631098, 12389, CellSignalling), mouse monoclonal GFAP antibody (1:200) (RRID: AB_2827276, SAB5201104, Sigma-Aldrich), mouse monoclonal CD31 (PECAM-1) (RRID: AB_10596359, BMS137, eBioscience), rabbit polyclonal CD31 (PECAM-1) (RRID: AB_10981955, PA5-16301, ThermoFisher), goat polyclonal PDGFRβ (RRID: AB_2162633, AF1042, R&D Systems), rabbit monoclonal PDGFRβ (RRID: AB_10985851, MA5-15143, ThermoFisher). The capillaries were washed twice with PBS, once in 0.5% BSA with 0.05% Tween20 in PBS for 10 minutes and incubated with secondary antibodies and Hoechst33342 (H1399, ThermoFisher) for 1 hour at RT in the dark. The following secondary antibodies were used: goat anti-mouse IgG (H+L) cross-adsorbed Alexa Fluor 546 (RRID: AB_2534071, A-11003, ThermoFisher), goat anti-mouse IgG (H+L) cross-adsorbed Alexa Fluor 488 (RRID: AB_2338840, 115-545-003-20, JacksonImmuno), goat anti-rabbit IgG (H+L) cross-adsorbed Alexa Fluor 488 (RRID: AB_2630356, ab150077, Abcam), donkey anti-goat IgG (H+L) cross-adsorbed Alexa Fluor 546 (RRID: AB_2534103, A11056, ThermoFisher)and Hoechst33342 (H1399, ThermoFisher). The capillaries were then washed three times in PBS and once in ddH_2_O for 10 minutes, air-dried, mounted with Keiser’s gelatine and imaged with Zeiss LSM880 confocal microscope.

### Primary mouse astrocyte culture and migration assay

Primary astrocytes were prepared from postnatal day 0 or 1 (P0/1) C57BL/6 mice. The cortical tissue was placed in a petri dish already containing a few drops of pre-warmed DMEM/F12 (10-090-CV, Corning) supplemented with 10% heat-inactivated foetal bovine serum (FBS) and 1% penicillin/streptomycin (p/s) (complete media). The tissue was cross-chopped with a scalpel and placed in a 15 ml falcon tube, also containing pre-warmed complete media. The tissue was then placed in the incubator for 15 minutes at 37°C. The tissue was gently triturated and the solution was passed through a 100 μm cell strainer (431752, Corning) before being gently centrifuged. The resulting pellet was re-suspended in complete media and plated in T75 flasks, one brain per flask. Cells were grown in the incubator at 37°C and 5% CO2 for 12-14 days and the media changed every 2-3 days. The grown cells were vigorously shaken for 2 hours to remove any contaminating microglial cells, trypsinized, resuspended in complete media and plated for experiments. Cells were used at passages 1 or 2. For migration experiments, astrocytes were serum-starved for 2 hours in T75 flasks, washed once with PBS and incubated with 0.25% trypsin for 10 minutes at 37°C. The media and free-floating astrocytes were then centrifuged, and the pellet was re-suspended in DMEM/F12 without FBS or p/s (serum-free media (SFM)) to get a 2.1 cells/ml cell suspension. The cells were plated in the top chamber of a transwell assay insert with an 8.0 μm transparent PET membrane (353097, Falcon). The top chamber contained 100 ng/ml LPS in SFM media (L4391, Sigma) with or without 1 μM clotrimazole (C6019, Sigma) and/or 10 μM NIBR189. Astrocytes were plated in SFM in the top chamber of the insert for 16-18 hours at 37°C, 5% CO2. The media inside the insert was discarded and the cells that didn’t migrate were removed with a cotton swab. The inserts were incubated in crystal violet solution (V5265, Sigma) for 10 minutes at room temperature (RT). Then, they were dipped carefully in a beaker with distilled water and left to air dry. When dried, images were taken with a light microscope at 20x magnification. Inserts were put in methanol (621990110, Poch) for 10 minutes on a shaker to extract the cells. The obtained solution was transferred to a 96-well plate and absorbance was read at 570/10 nm.

### Enzyme-linked immunosorbent assay (ELISA)

ELISAs were performed using DuoSet ELISA kits: mouse IL-6 (DY406-05), mouse IL-1b/1F2 (DY401-05) and mouse TNF-α (DY410-05) from R&D Systems according to the manufacturer’s instructions. Briefly, 100 μl of samples or standards in reagent diluent, 1% BSA in PBS, were added to a 96-well plate and incubated for 2 hours at RT. The wells were then emptied and washed three times with the wash buffer, 0.05% Tween-20 in PBS. Subsequently, 100 μl of the detection antibody diluted in the supplied reagent diluent was added to each well for 2 hours at RT. The plate was washed again with the wash buffer and 100 μl of streptavidin-HRP was added to each well for 20 minutes at RT in the dark. The plate was again washed three times and 100 μl of the substrate solution, 1:1 H_2_O_2_ and tetramethyl benzidine (34021, LifeTechnologies), was added to each well for 20 minutes at RT. To stop the reaction, 50 μl of H_2_SO_4_ 1 mol/L, was added to the wells and mixed gently. Optical density was read at 450 nm with a VICTOR Nivo plate reader (PerkinElmer).

### WB of the whole brain homogenates

The samples were equalised and mixed with sample buffer (1610747, Bio-Rad) + 10% β-mercaptoethanol, then incubated for 5min at 90 °C. The protein samples were separated on 10% SDS-polyacrylamide gels and transferred to PVDF membrane (IPFL00005, Merck) for semi-dry transfer. The membrane was blocked for 1h at RT in PBS with 0.1% Tween-20 (PBS-T), 5% non-fat milk and then incubated with primary antibody overnight at 4°C. The membrane was washed 3 times for 10 min and then incubated with a secondary antibody for 1h at RT. After 3 10-minute washes, the membrane was rinsed with water and read with WB scanner Odyssey. Band intensity was quantified using the software ImageJ and the gene of interest was normalised to the housekeeping gene. Primary antibodies used were: rabbit CH25H polyclonal antibody 1:1000 (600-401-MM8, Thermofisher), mouse CYP7B1 (OTI1G7) 1:500 (TA807549, Thermofisher), rabbit HSD3B7 1:1000 (RRID: AB_10856786, BS-2366R, Thermofisher), rabbit α-tubulin 1:5000 (RRID: AB_2546920, PA5-29444, Thermofisher) and mouse actin 1:5000 (RRID: AB_476744, A5441, Sigma) in PBS-T. The secondary antibodies used were goat anti-mouse IRDye 680RD (RRID: AB_621843, 925-32221, Li-Cor), donkey anti-mouse Alexa 790 (RRID: AB_2340870, 715-655-150, JacksonImmuno), goat anti-rabbit IRDye 800CW (RRID: AB_2651128, 925-68070, Li-Cor), goat anti-rabbit Alexa 680 (RRID: AB_2338085, 111-625-144, JacksonImmuno), 1:15000 in PBS-T.

### RNA sequencing

Gibco Human Astrocytes (cat. nr. K1884, Lot nr. 802268, Invitrogen) were sourced from a 20 weeks old male donor. No additional information was provided by the distributor. The Primary Human Brain Microvascular Endothelial Cells (cat. nr. ACBRI 376, Lot nr. RI-376, Cell Systems) were sourced from a 24 years old female donor. No other information is available. Human Brain Vascular Pericytes (cat. Nr. 1200, Lot nr. 10732, ScienCell) were sourced from a male donor. No other information is available. Conditionally immortalized human astrocytes clone 35 (HASTR/ci35), human brain vascular pericytes clone 37 (HBPC/ci37) and human brain microvascular endothelial cells (HBMECs) clone 18 (HBMEC/ci18) were then developed by professor Tomomi Furihata. RNA-sequencing was performed by Genewiz, Tokyo, Japan with 200 ng of total RNA from (i) untreated HBMECs, (ii) untreated or TNFα treated HBPCs; (iii) untreated or treated HASTR with TNFα/IL1α/C1q 50 ng/ml,10 ng/ml, 500 ng/ml respectively, all from PeproTech except for C1q from Calbiochem) or TNFα/IL1β for 96 hours. The NEBNext Ultra II RNA Library Prep Kit for Illumina (New England BioLabs, MA, USA) was performed on the RNA, after quality assessment, for cDNA library construction. Paired-end sequencing of 150 bp was used for RNA sequencing on the Illumina NovaSeq 6000 platform (Illumina, Inc., San Diego, CA, USA). The GRCh38 (hg38) was used to align the readings of the RNA sequence and quantification of gene expression levels was performed using HTSEQ v0.6.1. A different dataset from the same RNA-seq study was previously published: HBMECs (Ito et al. 2022) and HBPCs (Umehara et al. 2018).

### Data analysis

Five representative regions of z-stack images (maximum intensity projections), outside of the nuclei, were used for co-localisation analysis. The number of pixels present in each channel was obtained with the co-localisation tool of the Zen software. The number of co-localising pixels was normalised to the total number of pixels in the selected image (i.e. the sum of pixels in each separate channel plus the number of pixel present in both channels). The ratio of colocalised pixels was then averaged between the five representative images. The ratio ranged from 0, for no co-localisation, to 1, where the co-localisation is maximum. All statistical analysis was performed with GraphPad Prism 9 applying unpaired student t-tests for comparisons of two groups, one-way analysis of variance (ANOVA) for comparisons of three or more groups followed by Tukey’s multiple comparisons post-hoc tests or one-sample t-test for comparisons of three or more groups to normalised control. Data are shown as mean +/- standard error of the mean (SEM). Where appropriate, p values are written in the figure legends with significant effects indicated by asterisks: *p<0.05, **p<0.01, ***p<0.001, ****p<0.0001. Grubbs’ tests were run to identify and remove outliers.

## RESULTS

### The oxysterol 7α,25OHC synthesising (CH25H, CYP7B1) and degrading (HSD3B7) enzymes are present in the isolated mouse brain microvessels

In one of the first studies looking at the effects of LPS on oxysterol levels, mouse macrophages challenged with LPS upregulated the CH25H enzyme 35-fold and increased 6- to 7-fold their cellular concentration of oxysterol 25OHC(Diczfalusy et al. 2009). In another study, primary human monocyte-derived macrophages treated with LPS transiently increased expression of EBI2 and markedly upregulate the CH25H and CYP7B1 enzymes (Preuss et al. 2014). Similar upregulation of the CH25H and HSD3B7 enzymes was reported in LPS-treated BV2 microglia (Mutemberezi et al. 2018). We demonstrated that LPS-challenged primary mouse astrocytes synthesise and release oxysterols, including 7α,25OHC(Rutkowska et al. 2016). We also noted a rapid downregulation of the EBI2 transcripts upon LPS treatment and an increase in the mRNA levels of the CH25H and CYP7B1 enzymes. Moreover, the levels of the EBI2 ligand 7α,25OHC were highly increased in the media from LPS-treated astrocytes indicating these cells as one of the sources of oxysterols in the brain during inflammation. With the exception of CH25H which was recently shown to be synthesised by CNS ECs (Ruiz et al. 2023), the expression of EBI2 and the 7α,25OHC synthesising (CH25H, CYP7B1) and degrading (HSD3B7) enzymes in the brain microvascular cells has not been investigated. To examine the presence of EBI2 and the remaining enzymes in 7α,25OHC synthetic pathway in the brain vessels, we isolated normal mouse brain microvessels and quantified the receptor and enzyme levels. The data showed that EBI2 and the three enzymes were differentially present in the isolated normal mouse brain microvessels. EBI2 highly co-localised with pericytes (PGFRβ+ cells) and the ECs (CD31+ cells) and to a lesser degree with astrocytic endfeet (GFAP+) **(Figure 1B)**. The CH25H enzyme highly co-localised with CD31+ (ECs) and GFAP+ cells (astrocytic endfeet) and much less with PDGFRβ+ cells (pericytes) **(Figure 1C)**. The second enzyme in the 7α,25OHC synthetic pathway, CYP7B1, was moderately and to a similar degree present in CD31+ (ECs), GFAP+ (astrocytic endfeet) and PDGFRβ+ cells (pericytes) **(Figure 1D)**. The 7α,25OHC degrading enzyme HSD3B7 was highly present in CD31+ (ECs) and PDGFRβ+ cells (pericytes) and at very low levels in GFAP+ cells (astrocytic endfeet) **(Figure 1E)**.

### Systemic inflammation induces changes in the expression of *Ebi2* and *Ch25h, Cyp7b1* and *Hsd3b7* enzymes in the mouse brain

Peripheral injections of LPS were shown previously to induce neuroinflammation and disruption of the BBB (Banks et al. 2015; Li et al. 2020). We also demonstrated that repeated peripheral injections with LPS induce a significant increase in the number of pro-inflammatory cytokines in the mouse brain (Rutkowska et al. 2018). The disruption of the BBB function and the extent of neuroinflammation depend heavily on the applied protocol, particularly on the dose and frequency of LPS injections (Wang et al. 2014). In our study, we administered a single high dose (2 mg/kg) of LPS via intraperitoneal (i.p.) injection. This protocol effectively induced activation of astrocytes and microglia in the hippocampal region, along with the release of pro-inflammatory cytokines in the mouse brain. The cytokine release was observed at 12 hours (IL6 and IL1β) and 24 hours (IL1β) following the LPS injection **(Supplemental Figure 2)**. The protein and mRNA levels of *occludin* and *N-cadherin* were downregulated after 12 and 24 hours of LPS treatment **(Supplemental Figure 3)**.

In the EAE model, the concentration of the 7α,25OHC ligand in the CNS was sharply elevated as a result of the enhanced release of 7α,25OHC-synthesising enzymes, CH25H and CYP7B1, leading to an increased brain infiltration by peripheral immune cells (Wanke et al. 2017; Mutemberezi et al. 2018). Here, using the single high-dose LPS model, we investigated whether the EBI2/7α,25OHC system is affected in the brain during acute systemic inflammation **(Figure 2A)**. The *Ebi2* transcripts in whole brain homogenates were downregulated after 12 and 24 hours of LPS injection **(Figure 2B)**. The first 7α,25OHC synthesising enzyme, *Ch25h*, was strongly upregulated at both time points **(Figure 2C)** and *Cyp7b1* decreased 12 hours after the LPS challenge and then levelled with the vehicle-treated after 24 hours **(Figure 2D)**. The mRNA levels of the degrading enzyme, *Hsd3b7*, slowly increased after LPS treatment with significant changes observed after 24 hours **(Figure 2E)**. Quantification of protein levels of CH25H, CYP7B1 and HSD3B7 with WB showed similar trends but no statistically significant differences between the vehicle and LPS-treated mice **(Supplemental Figure 4)**.

**Fig. 2.**
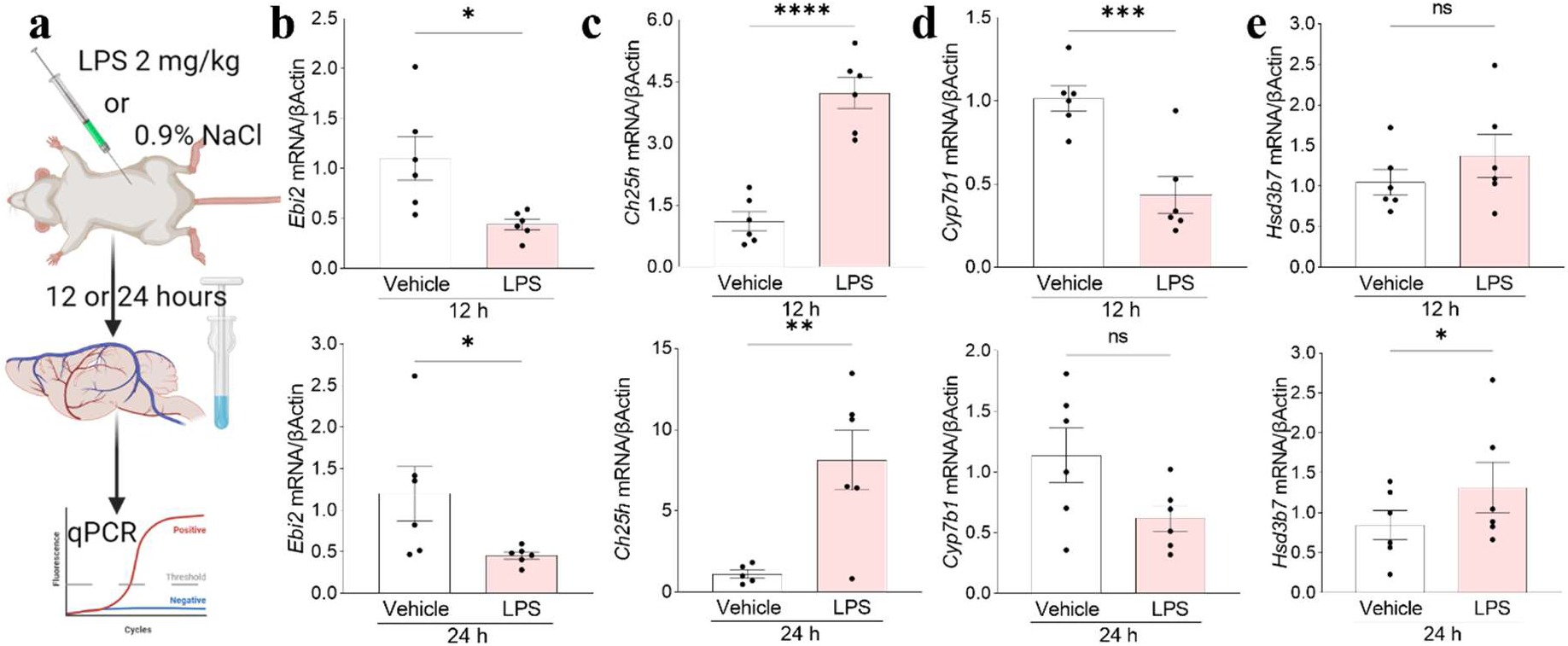
Systemic inflammation induces changes in the expression of Ebi2 and Ch25h, Cyp7b1 and Hsd3b7 enzymes in the mouse brain. a. Schematic representation of experimental setup. b. The expression of Ebi2 in whole brain homogenates decreased after 12 and 24 hours of LPS treatment (12 h: 40% +/- 2% vs. vehicle; 24 h: 38% +/- 4% vs. vehicle). c. The mRNA expression of the Ch25h enzyme in the mouse brain increased after 12 and 24 hours post LPS injection (12 h: 382% +/- 34% vs. vehicle; 24 h: 725% +/- 164% vs. vehicle). d. The mRNA levels of the Cyp7b1 enzyme decreased 12 hours after LPS treatment (43% +/- 11% vs. vehicle). e. The 7α,25OHC degrading enzyme, Hsd3b7, increased after 24 hours of LPS treatment (153% +/- 38% vs. vehicle). Data presented as mean + /− SEM, n=6 mice, unpaired student t-tests, *p<0.05, **p<0.01, ***p<0.001, ****p<0.0001 vs. corresponding vehicle.

### Systemic inflammation induces 7α,25OHC synthesising enzymes in the mouse brain vascular cells

After establishing that acute systemic inflammation modulates the expression of the EBI2/oxysterol system at the whole-brain level, we then set out to determine the contributions of the specific vascular cells. Increased levels of 7α,25OHC were also found *in vivo* in the spleens and brains of LPS-challenged mice (Hannedouche et al. 2011; Liu et al. 2011; Mutemberezi et al. 2018) and this LPS-mediated effect was not detected in CH25H knock-out mice (Hannedouche et al. 2011). Only recently, a study demonstrated that CNS ECs express CH25H, which, when ablated, attenuates EAE (Ruiz et al. 2023). Here, using the LPS model we investigated the changes in EBI2 and all enzymes involved in 7α,25OHC synthesis (Ch25H, CYP7B1) and degradation (HSD3B7). The increase in co-localisation of proteins after normalisation to the vehicle was quantified in immunohistochemically stained brain sections. The data indicated an increase in the number of co-localising pixels in EBI2+/GFAP+ and EBI2+/PDGFRβ+ cells and a smaller increase in EBI2+/CD31+ cells **(Figure 3A)**. The increase in CH25H enzyme co-localisation was observed for all three examined cell markers with a slightly greater mean increase in GFAP+ and PDGFRβ+ cells **(Figure 3B)**. The co-localisation of the CYP7B1 enzyme did not increase above the vehicle-treated levels in PDGFRβ+ cells but highly increased in CD31+ cells and negligibly in GFAP+ cells **(Figure 3C)**. The 7α,25OHC degrading enzyme, HSD3B7, was the least affected by the LPS challenge with only a slight increase in mean co-localisation in CD31+, GFAP+ and PDGFRβ+ cells **(Figure 3D)**. Because significant differences between human and mouse cells and various BBB models were observed and described before (Shawahna et al. 2013), we then examined the expression of the genes of interest in human brain microvascular endothelial cells (HBMECs), human brain vascular pericytes (HBPCs) and human astrocytes (HASTR) upon stimulation with various pro-inflammatory factors *in vitro*. HBMECs did not express EBI2, CH25H nor CYP7B1 but expressed the 7α,25OHC degrading enzyme HSD3B7. The expression of EBI2 in HBPCs was downregulated after stimulation with the pro-inflammatory cytokine TNFα and the expression of the first enzyme in the 7α,25OHC synthetic pathway, CH25H, was upregulated after stimulation. The second enzyme in the synthetic pathway, CYP7B1, was not expressed at baseline nor after stimulation with TNFα and the expression of the 7α,25OHC degrading enzyme, HSD3B7, was slightly upregulated after stimulation. HASTR expressed EBI2 and all three enzymes at baseline. Contrary to the downregulated expression observed in pericytes, EBI2 transcripts were induced after stimulation with pro-inflammatory cytokines. Similarly to HBPCs, the expression of CH25H was upregulated in stimulated HASTR. The expression of CYP7B1 remained low after stimulation and HSD3B7 was slightly downregulated after TNFα/IL1α/C1q and TNFα/IL1β stimulation. **(Supplemental Figure 5)**.

**Fig. 3.**
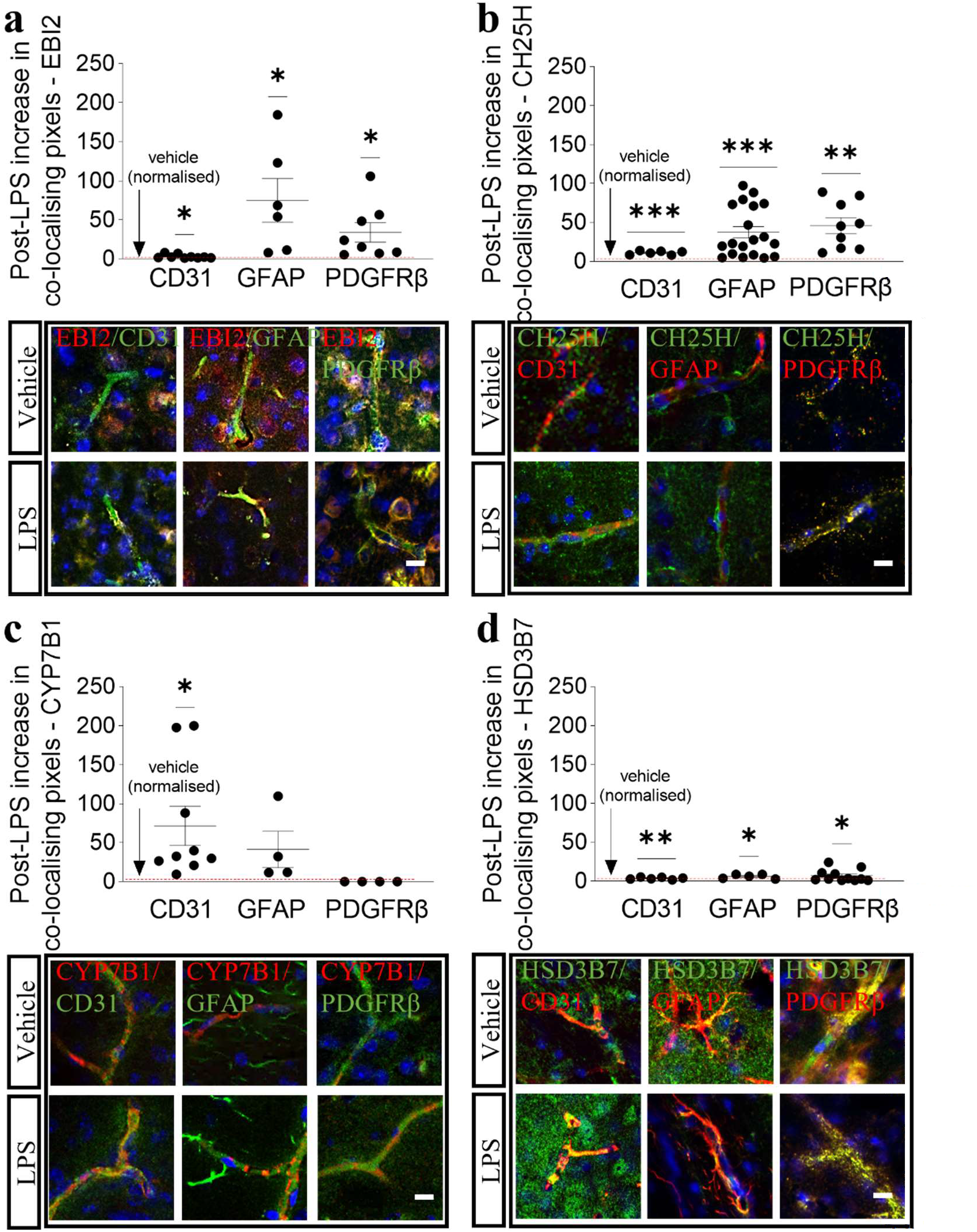
Systemic inflammation induces 7α,25OHC synthesising enzymes in the mouse brain vascular cells. a. The number of co-localising EBI2/CD31 pixels after LPS challenge increased onnly by 3.2% (+/- 0.8%). A moderate increase in the numer of co-localising pixels was recorded for EBI2/PDGFRβ (34% +/-12%) and EBI2/GFAP (75% +/-28%). b. After LPS challenge the numer of co-localising pixels of the first enzyme in the 7α,25OHC synthetic pathway, CH25H, with the EC marker, CD31, increased by 11% (+/-1%), with the pericyte/smooth muscle cel marker, PDGFRβ, by 46% (+/-10) and by 38% (+/-7.4%) with the astrocyte marker, GFAP. c. In contrast, the number of co-localising pixels of the second 7α,25OHC synthesising enzyme, CYP7B1, with CD31 increased by 72% (+/-25%), followed by a 41% increase (+/-23%) with GFAP positive cells and no change in PDGFRβ positive cells. d. The 7α,25OHC degrading enzyme, HSD3B7, was the least affected by systemic inflammation with the number of co-localising pixels increasing in CD31 positive cells by 3.8% (+/-0.56%), 7% (+/-2.4%) in PDGFRβ positive cells and 6.2% (+/-1.2%) in GFAP positive cells. Data presented as mean +/- SEM after normalisation to vehicle, one-sample t-test, *p<0.05, **p<0.01, ***p<0.001 vs. corresponding vehicle. Immunostaining of blood vessels in the cortex was performed on brain sections cut in the coronal plane. Scale 10 μm.

### The CYP7B1 inhibitor, clotrimazole, and the EBI2 receptor antagonist, NIBR189, attenuate LPS-induced astrocyte migration *in vitro*

EBI2 is a chemoattractant receptor shown to induce migration of EBI2-expressing CNS and immune cells *in vitro* and *in vivo* (Pereira et al. 2009; Gatto et al. 2009; Hannedouche et al. 2011; Preuss et al. 2014; Chalmin et al. 2015; Rutkowska et al. 2015; Clottu et al. 2017). Macrophages treated with media from LPS-treated astrocytes displayed increased migration, an effect inhibited with the EBI2 antagonist NIBR189, indicating EBI2/oxysterol-dependent mechanisms(Rutkowska et al. 2016). *In vivo* studies demonstrated that the trafficking of encephalitogenic CD4+ T cells into the CNS and the severity of EAE are significantly attenuated in CH25H-deficient mice (Chalmin et al. 2015). Increased CNS levels of 7α,25OHC during EAE were found to be the consequence of upregulated synthesis of CH25H by microglia and CYP7B1 by infiltrating lymphocytes and a simultaneous downregulation of HSD3B7. Clotrimazole, a CYP7B1 inhibitor **(Figure 4A)**, was shown *in vivo* to downregulate the levels of 7α,25OHC in the mouse spleen leading to altered B cell migration in the follicles, an effect similar to that observed in EBI2 KO mice (Liu et al. 2011). Here, we postulated that the concentration of EBI2 ligand increases in blood vessels localized at the inflammatory site, thereby inducing migration of peripheral and resident cells to the site of injury. To indirectly pre-test this hypothesis, we conducted *in vitro* experiments utilizing primary mouse astrocytes and we investigated the potential of clotrimazole and the EBI2 antagonist NIBR189 to inhibit the LPS-induced migration. The data showed increased migration of astrocytes upon LPS treatment **(Figure 4B)** and no increase in chemotaxis upon co-treatment with either clotrimazole or NIBR189, a selective EBI2 antagonist **(Figure 4C)**.

**Fig. 4.**
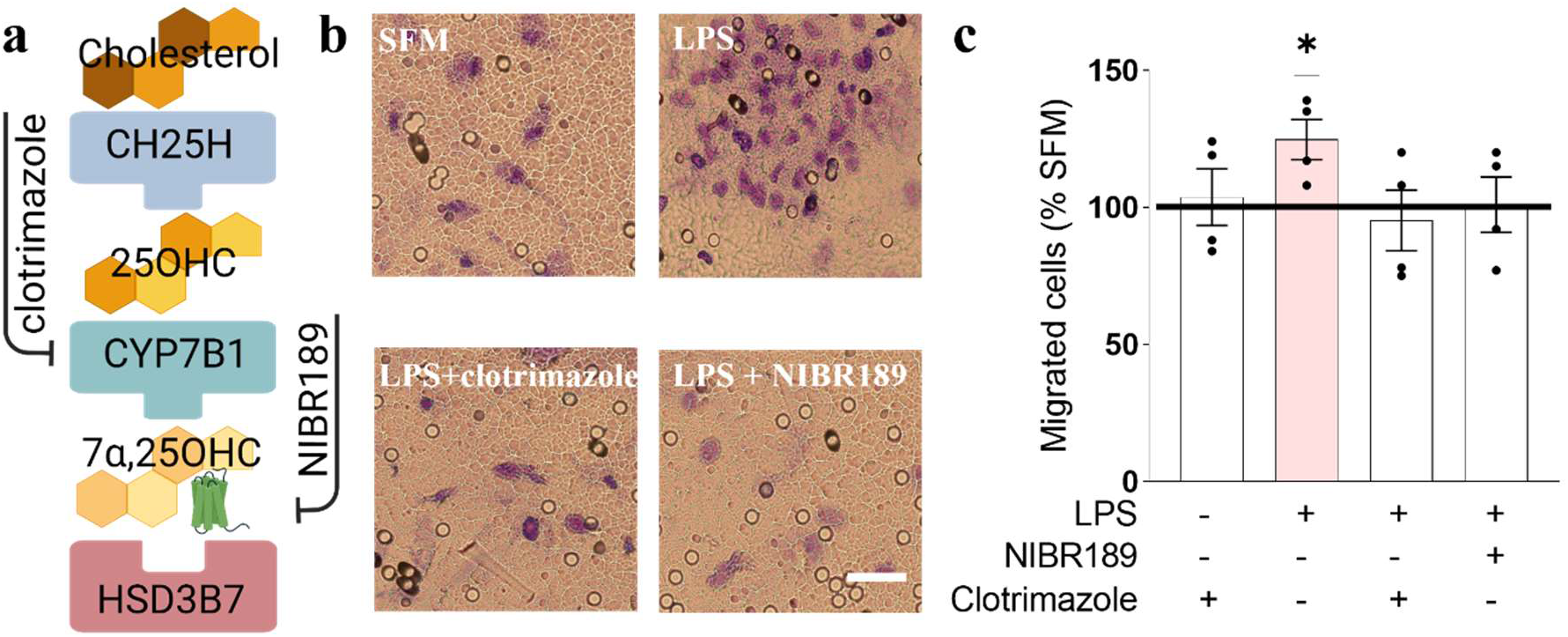
The CYP7B1 inhibitor, clotrimazole, and the EBI2 receptor antagonist, NIBR189, attenuate LPS-induced astrocyte migration in vitro. a. Schematic representation of 7α,25OHC synthetic pathway. b. Treatment of primary mouse astrocytes with LPS induces chemotaxis (125% +/- 7% vs. SFM). Data presented as mean +/− SEM, n=4 independent experiments, unpaired student t-test, *p<0.05. Scale 50 μm. c. Treatment of the mouse astrocytes with clotrimazole, a CYP7B1 inhibitor, or NIBR189, a selective EBI2 antagonist, decreased astrocyte migration (clotrimazole: 76% +/- 9% vs. LPS; NIBR189: 81% +/- 8% vs. LPS). Data presented as mean +/− SEM, n=4 independent experiments, one-sample t-test t(3)=3.366,*p<0.05. Scale 50 μm. SFM: serum-free media.

## DISCUSSION

The expression of EBI2 and the enzymes involved in 7α,25OHC synthesis and degradation is dynamically regulated under inflammatory conditions(Preuss et al. 2014; Chalmin et al. 2015; Rutkowska et al. 2016; Wanke et al. 2017; Crick et al. 2017; Mutemberezi et al. 2018). Identification of the cellular source of oxysterols in the brain, the magnitude and type of response (up-versus down-regulation), as well as the specificity of the cellular response to a given inflammatory factor, may help in the discovery of new drug targets for the modulation of immune cell trafficking into the CNS during systemic infection, neuroinflammatory or neurodegenerative diseases. Here, we first aimed to determine if the EBI2 receptor and CH25H, CYP7B1 and HSD3B7 enzymes are present in normal mouse brain microvessels. We established that EBI2 is highly present in ECs, pericytes/smooth muscle cells and astrocytic endfeet. The three enzymes (CH25H, CYP7B1 and HSD3B7) were also detected to a variable degree in each cell type indicating that the microvascular cells contribute to the synthesis, and possibly gradient formation and maintenance, of oxysterol 7α,25OHC in the mouse brain vasculature under normal conditions. We then determined if EBI2, CH25H, CYP7B1 and HSD3B7 are modulated in the brain vasculature under inflammatory conditions. Indeed, EBI2 and the 7α,25OHC synthesising enzymes, CH25H and CYP7B1, were highly modulated in the brain vasculature during acute inflammation. The degree of modulation differed depending on the enzyme and cell type indicating a specific microenvironment where levels of oxysterols are locally modulated and controlled. Moreover, differences between species and pro-inflammatory factors need to be taken into consideration when translating findings from animal models to humans. We here confirmed such differences between species in the levels of EBI2 and the three enzymes between mouse and human cells which form the BBB. EBI2 and the enzymes regulating its ligand levels are differentially expressed in the mouse and human vascular cells and are highly and distinctively modulated by different inflammatory factors. Such tightly controlled oxysterol microenvironment is also established and maintained in T cell follicles by differential CH25H, CYP7B1 and HSD3B7 enzyme expression by lymphoid stromal cells and follicular dendritic cells which locally increase or decrease 7α,25OHC levels, respectively (Yi et al. 2012). The established and maintained 7α,25OHC gradient is crucial for the appropriate localization of B cells in the lymphoid tissue and the launching of an immune response.

The concentration of 7α,25OHC in the mouse CNS increases in the EAE model of MS (Wanke et al. 2017; Mutemberezi et al. 2018). Again, the increase in 7α,25OHC levels results from differential regulation of CH25H, CYP7B1 and HSD3B7 enzymes by different cells including microglia and infiltrating lymphocytes (Wanke et al. 2017). The subsequent increase in the concentration of the EBI2 ligand, 7α,25OHC, was shown to enhance the migration of autoreactive T cells into the CNS and exacerbate the disease course. Similar results were obtained in the EAE model performed in CH25H knock-out mice where the disease severity was significantly attenuated as a result of reduced trafficking of memory CD4+ T cells into the CNS (Chalmin et al. 2015). Along the same lines, it was recently demonstrated that ablation of CH25H specifically in ECs attenuates EAE (Ruiz et al. 2023). Natalizumab, a disease-modifying therapy for MS, is a humanized IgG4 antibody against the α4-integrin subunit, which works by blocking the interaction between the α4 integrin expressed on lymphocytes and VCAM1 expressed on ECs thus limiting the passage of autoreactive lymphocytes across the vessel walls into the CNS (Brummer et al. 2021). Similarly, we demonstrated here that increased migration of EBI2-expressing cells, such as CNS resident astrocytes, in response to LPS treatment, is mediated by the EBI2/7α,25OHC system. The increase in chemotaxis was attenuated by inhibition of the ligand’s synthesis (clotrimazole) or directly by blocking the binding of the ligand with EBI2 (antagonist NIBR189). Modulation of the levels of CH25H, CYP7B1 and HSD3B7 enzymes directly in the brain microvasculature may thus be another way to limit the increased chemotaxis or entry of EBI2-expressing encephalitogenic immune cells into the CNS during acute inflammation, neuroinflammatory disease or neurodegenerative diseases.

At the whole brain level, we report downregulation of *Ebi2* transcripts after the LPS challenge while the EBI2 protein analysis indicated an increase of EBI2 in GFAP+ and PDFGRβ+ cells. Diverse cellular content in whole brain homogenates and different dynamics of EBI2 expression by various cells in response to inflammatory signalling may explain these contrasting observations. Indeed, different dynamics of EBI2 expression were reported before. For instance, we found a decrease in *Ebi2* mRNA levels in LPS-treated primary mouse astrocytes (Rutkowska et al. 2016), while others reported a temporary increase in *EBI2* expression in primary human macrophages after the LPS challenge (Preuss et al. 2014). Taken together, the combined *Ebi2* transcripts from all cell types present in the brain may indicate downregulation after LPS challenge while at the same time, specific cells may upregulate *Ebi2*. As a side note, endogenous downregulation of EBI2 during mycobacterium tuberculosis infection in macrophages was shown to contain early bacterial infection and intracellular survival (Tang et al. 2020). The downregulated *Ebi2* expression in the whole brain after the LPS challenge reported here may indicate a protective mechanism during bacterial infection (LPS challenge) in the CNS involving the EBI2 receptor.

Similarly, the expression of *Cyp7b1* initially decreased 12 hours after LPS treatment and started to return to baseline after 24 hours at the whole brain level. Whether the mRNA levels of *Cyp7b1* continued rising after 24 hours of LPS treatment remains to be elucidated, however, analysis of the protein levels, specifically in the brain microvascular cells indicated increased levels of CYP7B1 after LPS treatment in ECs and astrocytic endfeet and no induction in pericytes/smooth muscle cells (PDGFRβ+ cells). In a previous study of ours, we reported maximal mRNA expression of *Cyp7b1* after 24 hours of *in vitro* treatment with LPS while the levels of oxysterols 25OHC and 7α,25OHC peaked already after 15 hours indicating different dynamics of mRNA levels and oxysterol synthesis (Rutkowska et al. 2016). Moreover, Mutemberezi and colleagues (Mutemberezi et al. 2018) demonstrated that the LPS challenge induces 7α,25OHC levels in the mouse brain after as little as 4 hours and declines thereafter.

Lastly, the 7α,25OHC degrading enzyme, HSD3B7, was present in all three brain vascular cell types here studied and was least affected by inflammatory signalling, as was demonstrated by us before *in vitro* (Rutkowska et al. 2016). The RNA-seq of HBMECs, HBPCs and HASTR also demonstrated that the HSD3B7 enzyme is not affected by pro-inflammatory signalling. It is possible that during inflammation, the 7α,25OHC-synthesising enzymes, CH25H and CYP7B1, are upregulated and the degrading enzyme, HSD3B7, remains constant or is downregulated thus allowing for an increase in the concentration of 7α,25OHC in the inflamed tissue and a subsequent increase in immune cell infiltration of the inflamed tissue.

Taken together, these data indicate that the enzymes regulating the levels of 7α,25OHC are expressed directly by the brain microvascular cells. Moreover, the levels of EBI2 and CH25H, CYP7B1 are heavily regulated in the brain and the brain blood vascular cells during acute peripherally-induced inflammation. Modulation of EBI2 signalling and/or local concentrations of CH25H, CYP7B1 and HSD3B7 in the brain blood vessels might result in disease-modulatory effects with potential therapeutic applications in the treatment of neuroinflammatory diseases including multiple sclerosis.

## SUPPLEMENTARY FIGURES AND CAPTIONS

**Supp. Fig. 1.**
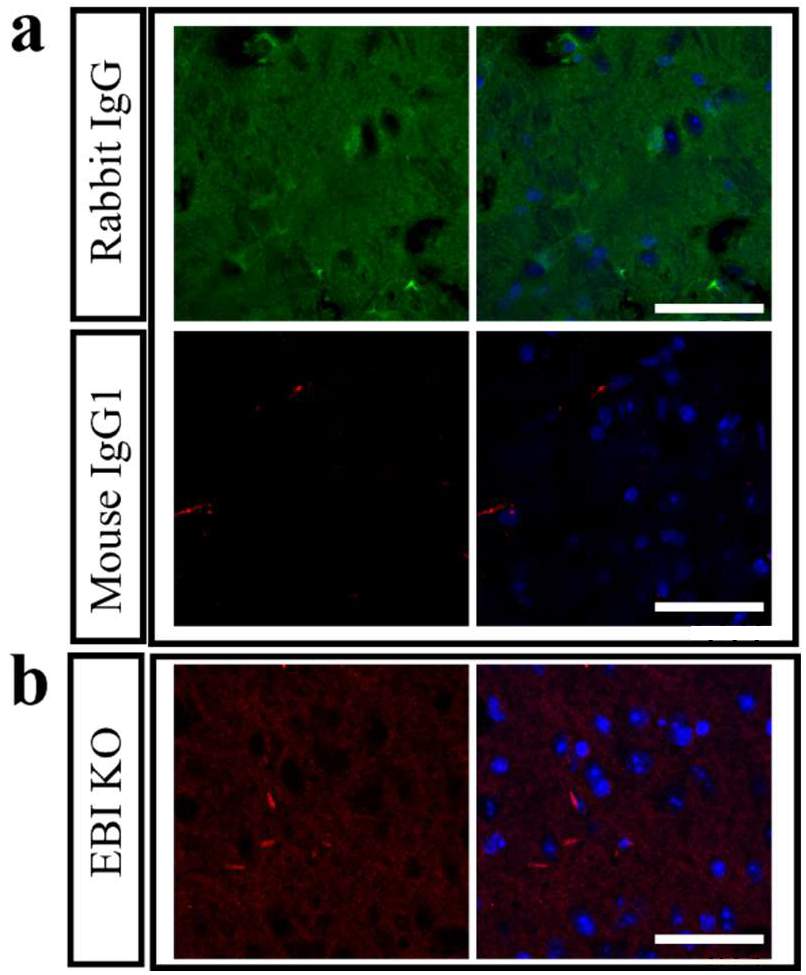
a. Representative images of mouse brain sections immunohistochemically stained with rabbit IgG and mouse IgG1 isotype control antibodies for rabbit anti-CH25H, rabbit anti-HSD3B7 and mouse anti-CYP7B1. B. Representative images of EBI2 knock-out (KO) brain sections stained for EBI2. Immunostaining of the cortex was performed on mouse brain sections cut in the coronal plane (sections from levels 70-87 Allen Mouse Brain Atlas). Scale 50 μm.

**Supp. Fig. 2.**
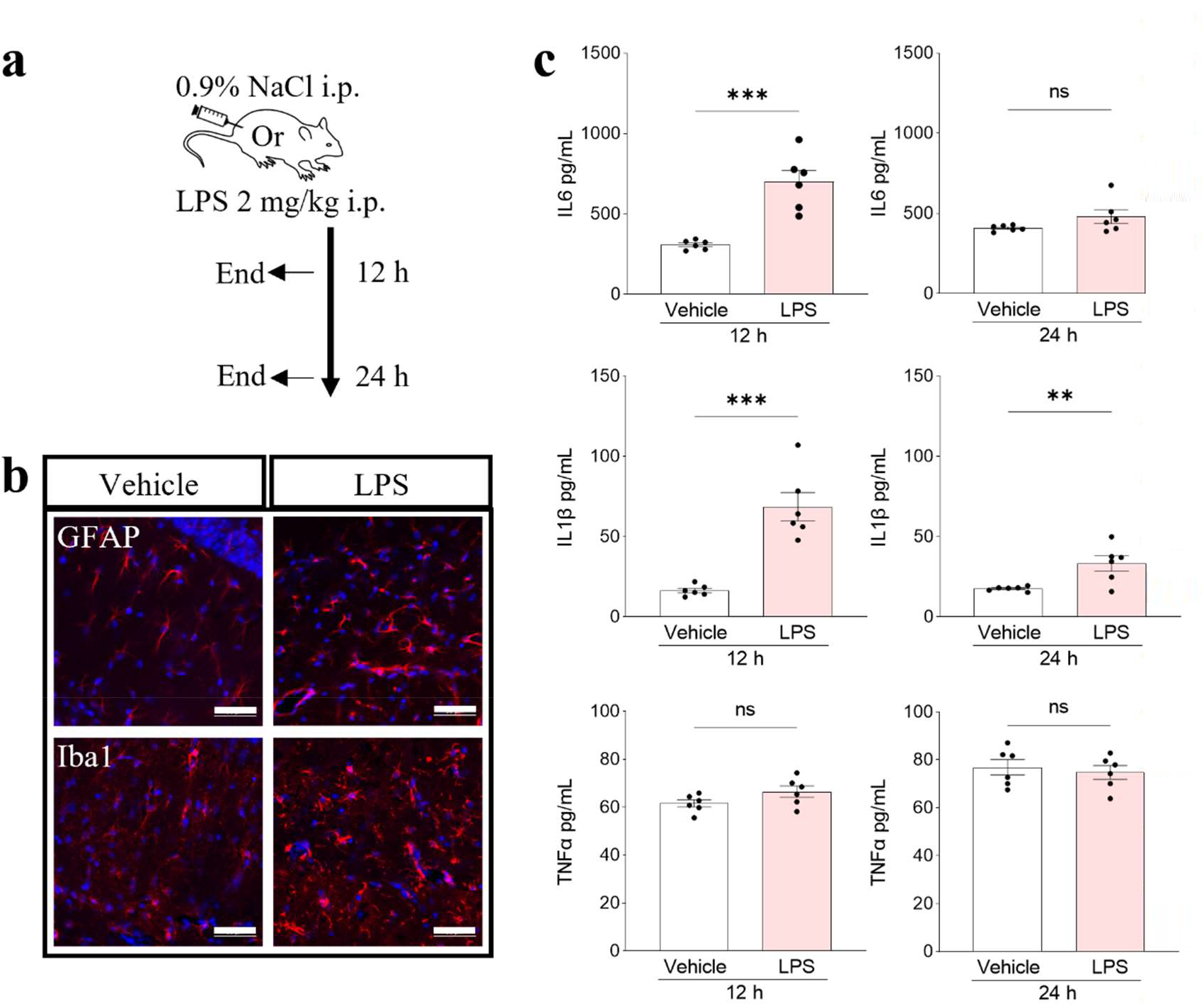
I.p. injection of LPS induces inflammation in the brain. a. Schematic representation of the experimental protocol. b. I.p. injection of LPS induces mild neuroinflammation as indicated by increased astrocyte (GFAP, red) and microglia (Iba1, red) reactivity. Representative images, scale 50 μm. Immunostaining was performed on mouse brain sections cut in the coronal plane (sections corresponding to levels 70-87 in Allen Mouse Brain Atlas). Images show GFAP and Iba-positive cells in the hippocampal region. c. The levels of pro-inflammatory cytokines in the whole brain homogenates increased after 12 h (IL6: 227% +/- 23% vs. vehicle; IL1β: 416% +/- 53% vs. vehicle) and 24 h (IL1β: 189% +/- 27% vs. vehicle). TNFα levels did not change after LPS treatment. Data presented as mean + /− SEM, n=6 mice, unpaired t-test, **p<0.01; ***p<0.001 vs. corresponding vehicle.

**Supp. Fig. 3.**
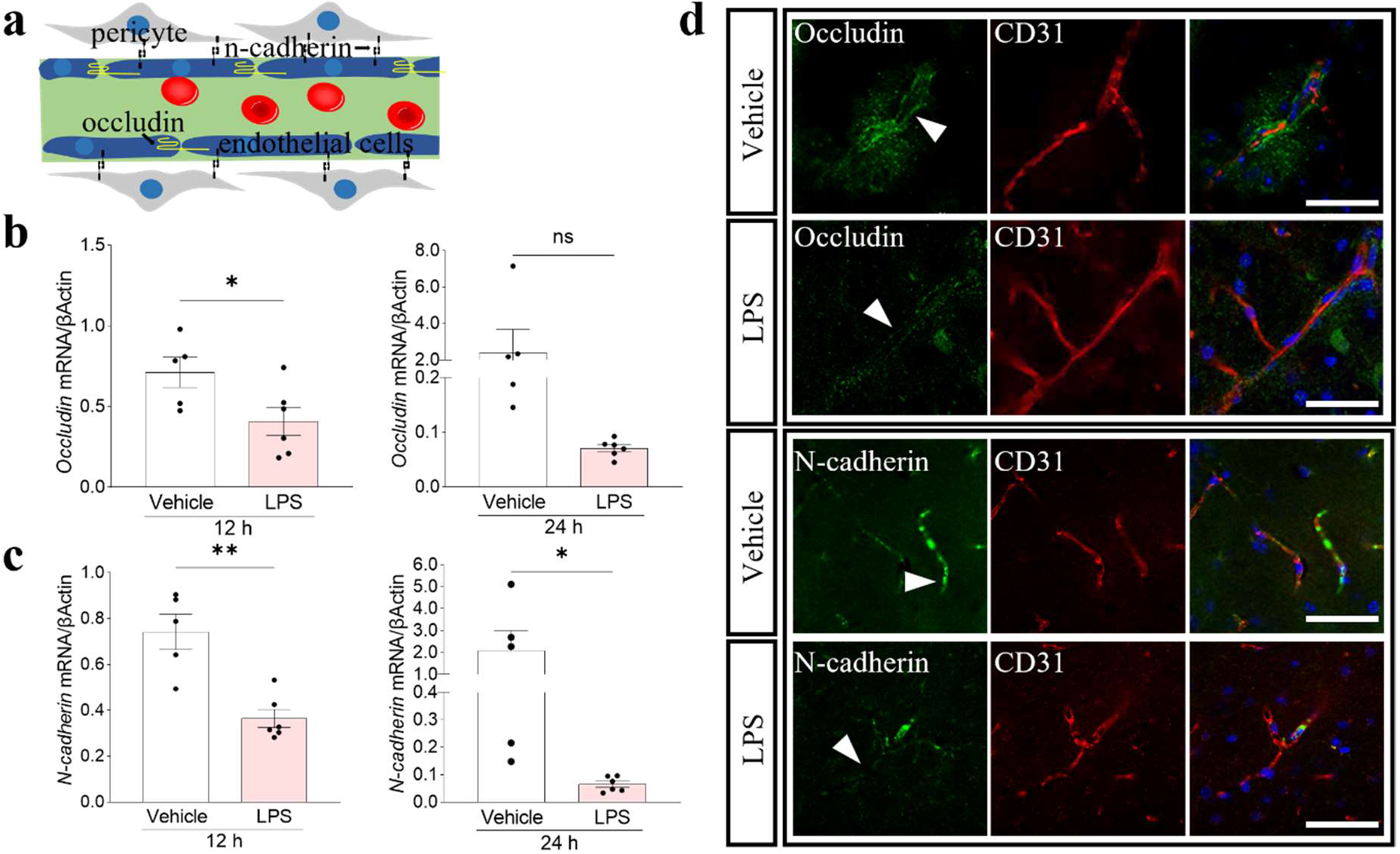
Tight junction and adhesion proteins in the brain are downregulated during systemic inflammation. a. Simplified graphical representation of the BBB showing the localisation of occludin (yellow) forming tight junctions between ECs, and the localisation of N-cadherin (black), an adhesion protein between the pericytes and ECs. b. The mRNA levels of occludin were significantly reduced after 12 hours post LPS injection (12 h: 57% +/- 12% vs. vehicle) and started to return to the vehicle levels after 24 hours. c. Expression of the adhesion molecule N-cadherin significantly decreased after 12 hours (12 h: 49% +/- 5% vs. vehicle) and 24 h (3% +/- 1% vs. vehicle) of LPS treatment. Data presented as mean + /− SEM, n=5-6 mice, unpaired t-test, *p<0.05, **p<0.01 vs. corresponding vehicle. d. Representative confocal microscope images show decreased occludin (green) and N-cadherin (green) proteins after LPS treatment in the mouse brain blood vessels (ECs, CD31, red). Immunostaining of cortical blood vessels was performed on mouse brain sections cut in the coronal plane (sections corresponding to levels 70-87 in Allen Mouse Brain Atlas). Scale 50 μm.

**Supp. Fig. 4.**
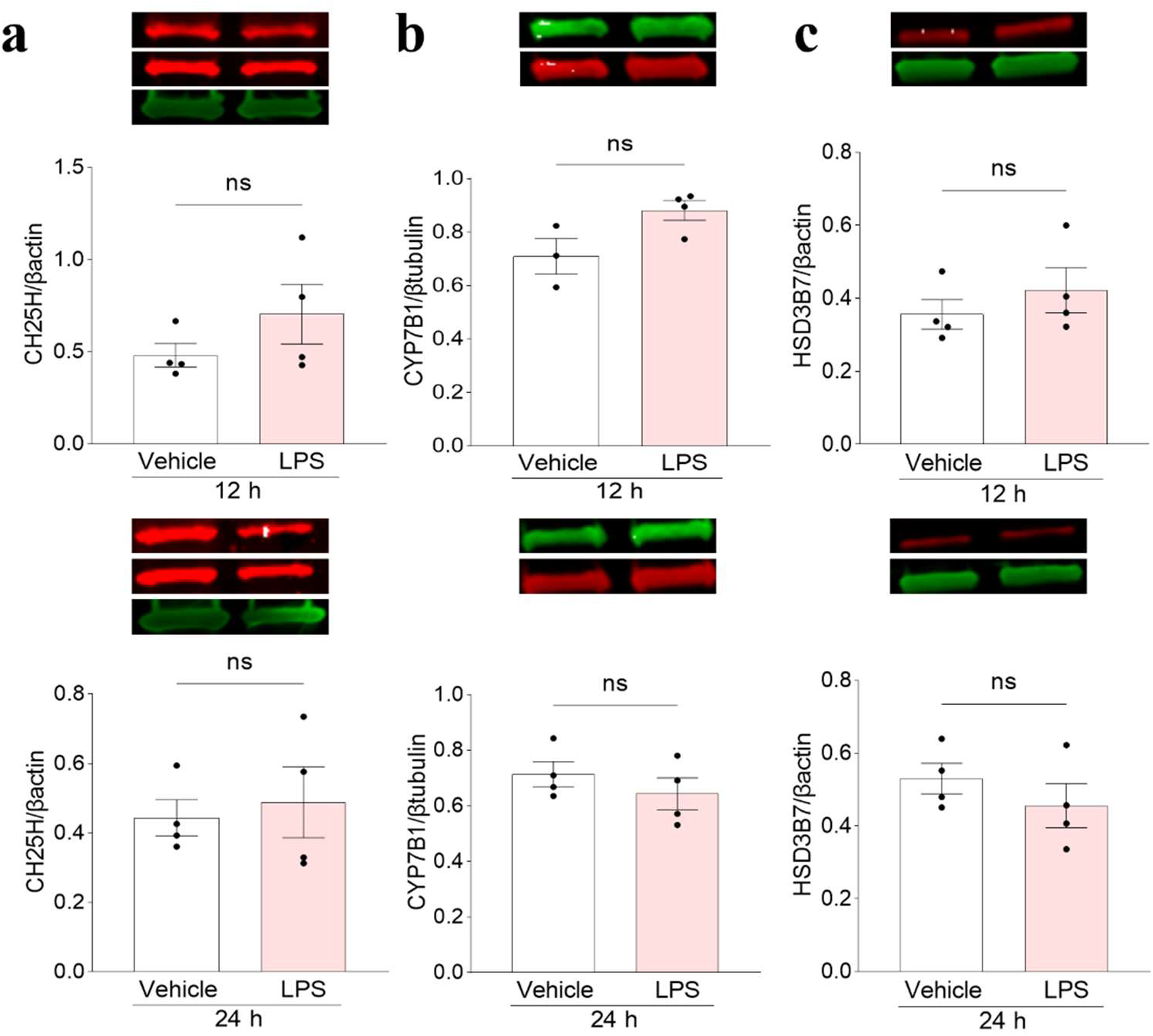
a-d. Quantification of CH25H, CYP7B1 and HSD3B7 protein levels in whole mouse brain homogenates. Data presented as mean +/- SEM, unpaired student t-tests.

**Supp. Fig. 5.**
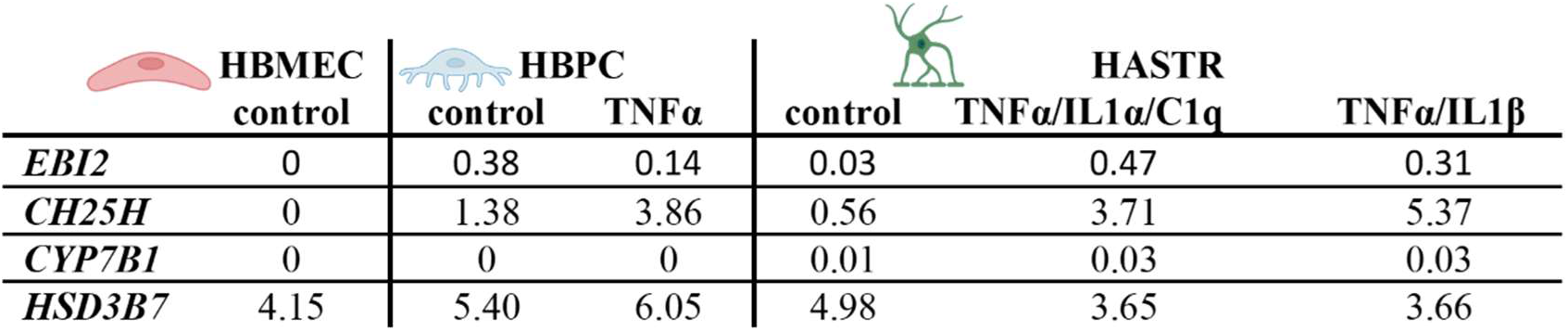
Expression of EBI2, CH25H, CYP7B1 and HSD3B7 enzymes in human brain vascular cells. A. Single-cell RNA-seq data showing expression of EBI2, CH25H, CYP7B1 and HSD3B7 in human brain microvascular ECs (HBMECs), human brain vascular pericytes (HBPCs) and human brain astrocytes (HASTRs) at baseline (unstimulated/control cells) and after stimulation with TNFα (HBPCs), TNFα/IL1α/C1q or TNFα/IL1β (HASTRs). Gene expression unit is: fragments per kilobase of transcript per million mapped fragments (FPKM).

## STATEMENTS AND DECLARATIONS

### Author’s contributions

A.R. conceptualised the study, acquired funding, supervised the study and did the visualisation. B.K. and A.R. are responsible for the project administration and coordination. F.C. and A.R. designed the methodology, performed the investigations, data analysis and wrote the original draft. I.K. performed the mouse *in vivo* model. T.F. performed the RNA-seq study, developed HASTR, HBPC, HBMEC cells and acquired funding for RNA-seq analysis. B.K and T.F. reviewed and edited the manuscript. All authors reviewed and accepted the final version of the manuscript.

### Data availability

The data is available on public repository: https://zenodo.org/record/8330841

### Ethics approval and consent to participate

All animal experiments were approved by the Local Ethical Committee for Animal Experiments in Bydgoszcz, Poland under licence numbers 27/2019 and 38/2021.

## Acknowledgements

We would like to thank Ms Beata Muszyńska from the Tricity Academic Experimental Animal House for her support with the *in vivo* mouse model.

